# CYLD, a Mechanosensitive Deubiquitinase, Regulates TGFβ Signaling in Load-Induced Bone Formation

**DOI:** 10.1101/665802

**Authors:** Jacqueline Nguyen, Ramin Massoumi, Tamara Alliston

## Abstract

Many signaling pathways involved in bone homeostasis also participate in the anabolic response of bone to mechanical loading. For example, TGFβ signaling coordinates the maintenance of bone mass and bone quality through its effects on osteoblasts, osteoclasts, and osteocytes. TGFβ signaling is also essential for the mechanosensitive formation of new bone. However, the mechanosensitive mechanisms controlling TGFβ signaling in osteocytes remain to be determined, particularly those that integrate TGFβ signaling with other early responses to mechanical stimulation. Here, we used an in vivo mouse hindlimb loading model to identify mechanosensitive molecules in the TGFβ pathway, and MLOY4 cells to evaluate their interactions with the prostaglandin E2 (PGE2) pathway, which is well-known for its rapid response to mechanical stimulation and its role in bone anabolism. Although mRNA levels for several TGFβ ligands, receptors, and effectors were unchanged, the level of phosphorylated Smad3 (pSmad3) was reduced in tibial bone as early as 3 hrs after early mechanical stimulation. We found that PGE2 and its receptor, EP2, repress pSmad3 levels and transactivation of Serpine1 in osteocytes. PGE2 and EP2 control the level of pSmad3 through a proteasome-dependent mechanism that relies on the deubiquitinase CYLD. CYLD protein levels were also reduced in the tibiae within 3 hrs of mechanical loading. Using CYLD-deficient mice, we found that CYLD is required for the rapid load-mediated repression of pSmad3 and for load-induced bone formation. These data introduce CYLD as a mechanosensitive deubiquitinase that participates in the PGE2-dependent repression of TGFβ signaling in osteocytes.

## 1. Introduction

Mechanical loading on bone is required for the maintenance of bone mass. In astronauts, sedentary adults, or in mice with unloaded hindlimbs, the reduced mechanical stimulation deregulates mineral homeostasis and causes bone loss [1] [2] [3]. Since low bone mass diseases such as osteoporosis predispose 125 million people worldwide to the risk of fractures [4], understanding the cellular and molecular mechanisms by which mechanical load maintains bone homeostasis is clinically significant. Osteocytes are the bone-embedded cells that sense mechanical stimuli and respond by regulating osteoblast and osteoclast activity [5]. For example, mechanical loading represses the expression of an osteocyte-secreted protein, called Sclerostin, which binds and inhibits the Wnt coreceptor, Lrp5/6 on osteoblasts. Thus, loading stimulates Wnt-mediated bone anabolism by repressing Sclerostin levels [6] [7].

Previously, we found that mice with reduced TGFβ signaling in osteocytes cannot relay the load-mediated repression of Sclerostin expression and, consequently, have impaired load-induced bone formation [8]. The mechanoregulation of Sclerostin expression and new bone formation required the active phosphorylated form of the TGFβ effector Smad3, which was repressed within 5 hrs of loading. However, the mechanosensitive mechanisms regulating TGFβ signaling in osteocytes remain unclear, particularly those that occur rapidly in response to loading. Here we sought to elucidate early events in the anabolic response of bone to mechanical load by focusing on the mechanisms responsible for the repression of phosphorylated Smad3 within 5h of mechanical stimulation.

A rapid increase in prostaglandin E2, or PGE2, is one of the first responses of osteocytes to mechanical stimulation. Through a variety of mechanisms, PGE2 leads to increased transcription of genes that elicit bone formation. PGE2 facilitates cell-cell communication by moving between cells through gap channels, like connexin43, to maintain bone homeostasis [9]. Fluid flow on osteocytes induces connexin43 translocation to the cell membrane, allowing for PGE2 to move into the adjacent cell [10]. PGE2 also acts extracellularly by binding to transmembrane receptors. PGE2 binding to the EP2 receptor in osteocytes stimulates PKA and PI3K signaling to repress the antagonist of Wnt/β-catenin signaling, GSK3β [11] [12]. PGE2 further induces Wnt signaling by repressing Sclerostin expression [13]. In addition, PGE2 produced from mechanically stimulated osteocytes protects osteocytes from glucocorticoid-induced apoptosis [14]. These are some of the mechanisms by which PGE2 rapidly coordinates the activity of multiple signaling pathways to maintain osteocyte function and bone homeostasis in response to mechanical load.

Although crosstalk between PGE2 and TGFβ signaling is not defined in osteocytes, in other tissues PGE2 plays an important role in the control of matrix deposition and TGFβ signaling. For example, PGE2 represses TGFβ through an EP2/PKA-dependent mechanism to stimulate fibroblast activity [15]. PGE2-activated EP4 signaling in prostate cancer cells relies on TGFβ signaling to stimulate cell migration and metastasis [16]. PGE2-activated EP2 regulates TGFβ-induced mammary fibrosis and oncogenesis [17]. Since TGFβ mediates PGE2 effects in multiple biological systems, we hypothesized that PGE2 regulates TGFβ signaling in osteocytes, possibly as part of a signaling cascade that occurs rapidly after mechanical stimulation.

Among the possible mechanisms to rapidly reduce the level of signaling molecules is ubiquitin-dependent proteasomal degradation. In other cell types, this process has been implicated as a mechanosensitive mechanism. In muscle cells, the ubiquitin ligase, CHIP, marks damaged proteins for degradation when cells are subjected to tensional pull [18]. *C. elegans* utilizes the E3 ubiquitin ligase, MTB1, to direct the degradation of a mechanosensitive channel called MEC4, which allows the organism to sense touch [19]. However, mechanosensitive control of ubiquitin-dependent proteosomal activity in osteocytes has not been evaluated.

Ubiquitin is also an important regulatory mechanism in TGFβ signaling and in bone homeostasis. In particular, CYLD is a deubiquitinase that indirectly targets Smad3 for degradation in lung cells to regulate matrix deposition and fibrosis [20]. In addition to regulating the TGFβ pathway, CYLD targets NEMO, a subunit of IKK, and represses NF-kB signaling [21][22]. The NF-kB pathway is important for regulating RANK signaling in osteoclastogenesis. CYLD deficient mice have increased osteoclast numbers and activity and, consequently, low trabecular bone mass [23]. Since Smad3 is rapidly downregulated in mechanically-stimulated bone, we hypothesized that CYLD participates in load-mediated repression of TGFβ signaling in osteocytes and in load-induced bone formation.

Although the importance of TGFβ in osteocyte mechanotransduction is established, the mechanosensitive mechanisms controlling TGFβ signaling in bone, particularly its interaction with PGE2, remain to be determined. The regulated turnover of signaling complexes is an essential mechanism for the control of signal transduction and cellular function. However, the role of ubiquitin-dependent proteasomal degradation in osteocyte mechanotransduction is not well-studied. Therefore, we investigated mechanosensitive mechanisms controlling Smad3 phosphorylation and load-induced bone formation, with a focus on PGE2 and the deubiquitinase CYLD.

## 2. Materials and Methods

### 2.1 Mice

Treatments and protocols used for the animal studies were approved by the University of California, San Francisco Institution Animal Care and Use Committee and were designed to minimize discomfort to the animals. This study used 8-9 week-old male CYLD-deficient mice on a C57BL/6 background [24]. Wild type littermates were used as comparative controls.

### 2.2 In Vivo Mechanical Loading

Axial compressive loads equivalent to 10 times the mouse’s body weight were delivered by a Bose Electroforce ELF3200 desktop load frame (Bose, MN, USA) fitted with two custom-made hemi-spherical fixtures that gripped the mouse knee and ankle [8]. Similar methods of in vivo loading have been shown to upregulate bone anabolism [25]. In our analysis of these ex vivo limb loading conditions using in situ strain rosettes, this range of parameters produces maximum principal strains in the range of 1500-2500 με on the mid-diaphyseal surface of the tibiae. For each mouse, only the right hind limb was loaded, while the left hind limb was not loaded to serve as the contralateral control (nonloaded). Each round of loading consisted of 600 cycles of axial compression at 1 Hz for 10 minutes administered under general injectable anesthesia (ketamine). The anesthesia recovery agent, atipamezole, was not used. Unless otherwise indicated, mice were subjected to this loading regimen once and were euthanized at the indicated time points after loading. For analysis of load-induced bone formation in CYLD-deficient mice, mice were loaded for 5 consecutive days as described below.

### 2.3 Histomorphometry

Mice were loaded for 5 consecutive days (with one bout of loading each day) with calcein injected at day 1 and day 5 of the loading regimen. Two days after the last loading, mice were euthanized and the tibiae were collected and stored in 4% paraformaldehyde in PBS. Bones were processed and embedded in methyl methacrylate for sectioning to generate 5 μm sections for analysis of calcein labeling with an epifluorescent microscope at 20X magnification. Calcein labels were measured using ImageJ.

### 2.4 Cell Culture

MLO-Y4 cells (gift from Dr. Lynda Bonewald), a mature mouse osteocyte cell line, were used for the in vitro studies. MLO-Y4 cells were cultured as described on plastic tissue culture plates coated with rat-tail collagen I (BD). These cells are cultured in 2.5% heat-inactivated Fetal Bovine Serum (Characterized, Hyclone) and 2.5% heat-inactivated Bovine Calf Serum (UCSF CCF) in alpha-MEM media containing nucleosides (UCSF CCF). At 80% confluence, MLO-Y4 cells were treated with the following reagents in serum-free media: 5 ng/ml TGFβ1 (Peprotech, 100-36E), 0.1 μM PGE2 (Sigma), 1 μM Butaprost (EP2 agonist, Cayman), 50uM MG132 (proteasome inhibitor, Sigma), 20uM AH6809 (EP2 antagonist, Cayman), or 20uM PGE1 alcohol (EP4 agonist, Cayman) for the indicated times. All agents were resuspended in solvents suggested by product manufacturers. For controls, vehicle was added to cells in the same conditions.

### 2.5 siRNA or shRNA knockdown of CYLD

CYLD knockdown was achieved using a stably expressed lentiviral pLKO.1 puro mouse CYLD shRNA construct (Sigma, MC5) and nontarget shRNA (Sigma, SHC002). Lentiviral vectors were packaged into the virus by the UCSF RNAi Core. MLOY4 cells were grown to 80% confluency and infected with CYLD or nontarget lentivirus. MLOY4 cells with the shRNA vectors were selected with 5 μM Puromycin and maintained under selection throughout the experiments. We were able to achieve about a 50% knockdown verified through qPCR of the nontarget and CYLD shRNA transfected cells.

CYLD knockdown using siRNA was achieved using mouse CYLD siRNA (AUAACUAGAUACGUAAGACUU) and scrambled (GUCUUACGUAUCUAGUUAUUU). siRNA oligos were transfected into MLOY4 cells using Lipofectamine2000. Cells were treated as indicated 48-72 hrs after transfection. RNA and protein were extracted as indicated.

### 2.6 Quantitative PCR

RNA from the tibia bone was extracted by first removing the epiphyses, and subsequently the bone was centrifuged to remove the marrow. The bones were snap frozen in liquid nitrogen and pulverized with a mortar and pestle. Bone powder was transferred to Trizol (Invitrogen) and RNA was purified using the Invitrogen Purelink system according to the manufacturer’s instructions. RNA was extracted from cells using the Qiagen RNeasy Mini Kit following the manufacturer’s instructions. RNA concentration was measured using a Nanodrop ND1000 Spectrophotometer. cDNA was generated from up to 1 μg of RNA with the iScript cDNA Synthesis kit (Bio-Rad). Quantitative PCR was performed using iQ SYBR Green Supermix (Bio-Rad) with the primers on the table below. Each gene was normalized against the housekeeping gene, L19. Analysis used the ΔΔCT method to compare the load vs nonloaded samples from the same animal.

**Table 2.1:**
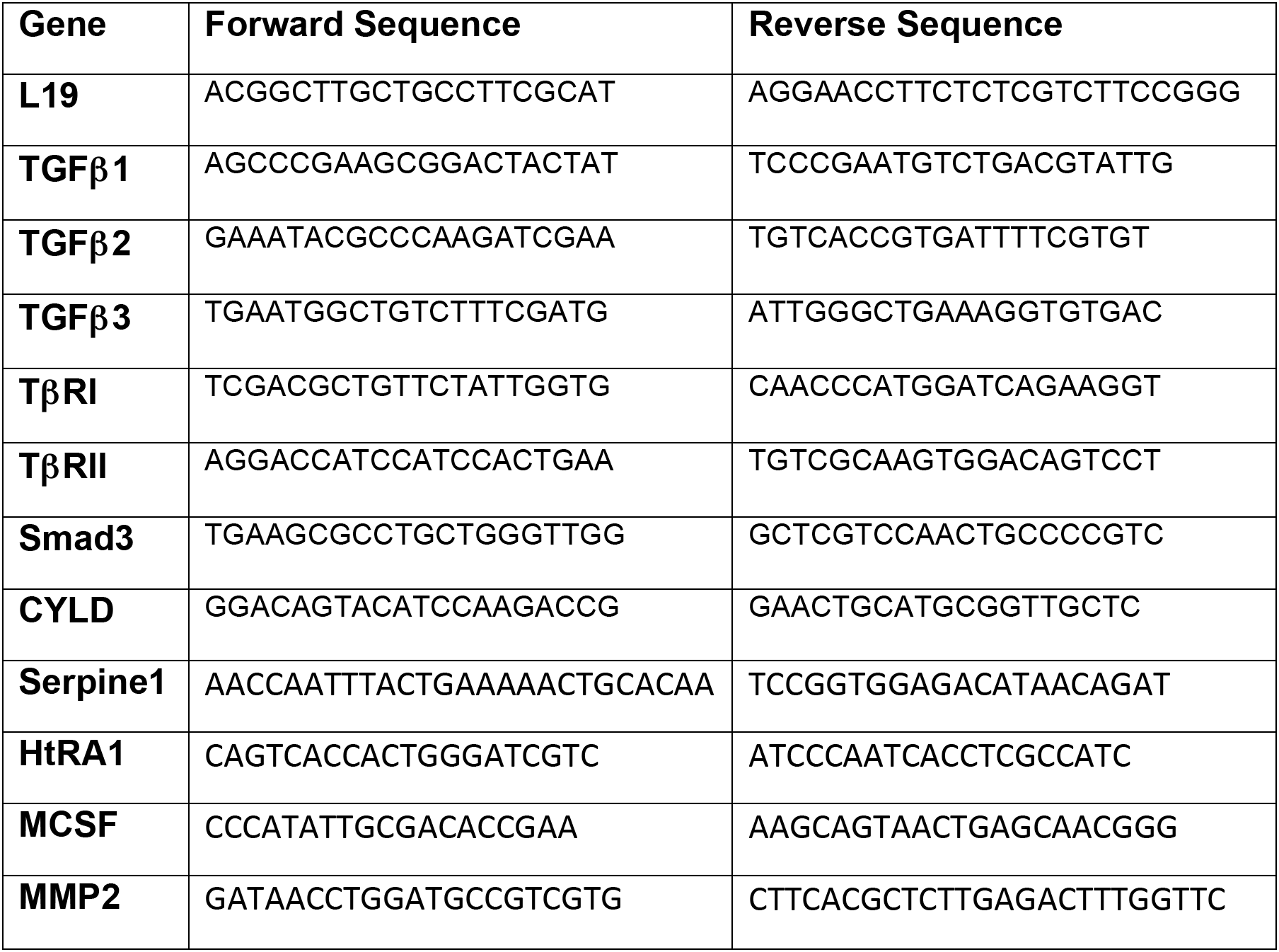
Primers for QPCR of genes of interests.

### 2.7 Western Blot Analysis

Bone protein was isolated from liquid nitrogen snap frozen tibia by pulverizing the bone using a mortar and pestle and transferring the powder to radioimmunoprecipitation buffer (RIPA) containing 10 mM Tris, pH 8, 1 mM EDTA, 1 mM EGTA, 140 mM sodium chloride, 1% sodium pyrophosphate, 100 mM sodium fluoride, 500 μM PMSF, and 5 mg/mL eComplete Mini protease inhibitor tablet (Roche). Lysates were sonicated for 10 seconds 8 times in an ice-cold water bath and centrifuged to separate the protein extract. Bradford assay was used to quantify the amount of proteins and similar amount of proteins were loaded onto the gel. The protein supernatant was loaded onto a 10% SDS-PAGE gel, separated, and transferred to a nitrocellulose membrane. Similarly, MLO-Y4 whole cell lysate was collected in RIPA buffer, sonicated for 10 seconds 3 times in ice-cold water bath, for separation on a 10% SDS-PAGE as above. Westerns were probed using the following antibodies: mouse anti-beta actin (Abcam), rabbit anti-pSmad3 (gift from Dr. E. Leof), rabbit anti-Smad2/3 (Santa Cruz Biotech or abcam), rabbit anti-TβRI (Santa Cruz Biotech), and rabbit anti-CYLD (Sigma). Fluorescently-tagged secondary antibodies were used to detect the primary antibodies above and the blots were imaged on an Odyssey LiCor Imaging System. The protein bands were normalized to the housekeeping protein, beta-actin, and the intensity was measured on the LiCor imaging software according to manufacture’s instructions.

### 2.8 Statistical Analyses

Statistics were performed using GraphPad Prism 5. For the animal studies, at least 5 animals were used for each analysis. For the in vitro studies, each experiment had two technical replicates and was repeated at least three times. T-tests were used to compare the differences between groups of normally distributed data. Nonparametric tests were used for non-normally distributed data as stated in each figure. Significance of comparisons is defined by p-values equal to or less than 0.05.

## 3. Results

### 3.1 Rapid post-transcriptional control of TGFβ mechanosensitivity

We previously reported the rapid mechanosensitive repression of TGFβ signaling within 5 hrs of in vivo hindlimb loading [8]. Although the mechanoregulation of TGFβ signaling was essential for load-induced bone formation, the mechanisms responsible for TGFβ mechanosensitivity in bone remain unclear. To elucidate these mechanisms, we examined the effect of short-term in vivo hindlimb loading on components of the TGFβ signaling pathway, including TGFβ ligands, receptors, Smads, and other known TGFβ effectors. Consistent with our prior findings at 5 hrs after hindlimb loading [8], the levels of Smad3 phosphorylation were decreased within 3 hrs of load, relative to nonloaded tibiae (Figure 1A). TβRI protein levels were also reduced with load, but the protein reduction was not statistically significant as compared to pSmad3 levels. However, transcriptional analysis of mRNA encoding TGFβ ligands (TGFβ1, TGFβ2, TGFβ3), receptors (TβRI and TβRII), and Smads (Smad3) show no significant differences between nonloaded and loaded tibiae (Figure 1B). Other factors implicated in the post-transcriptional regulation of TGFβ signaling, including HtRA1 and CYLD, also show no load-dependent differences in mRNA expression (Figure 1B).

**Figure 1:**
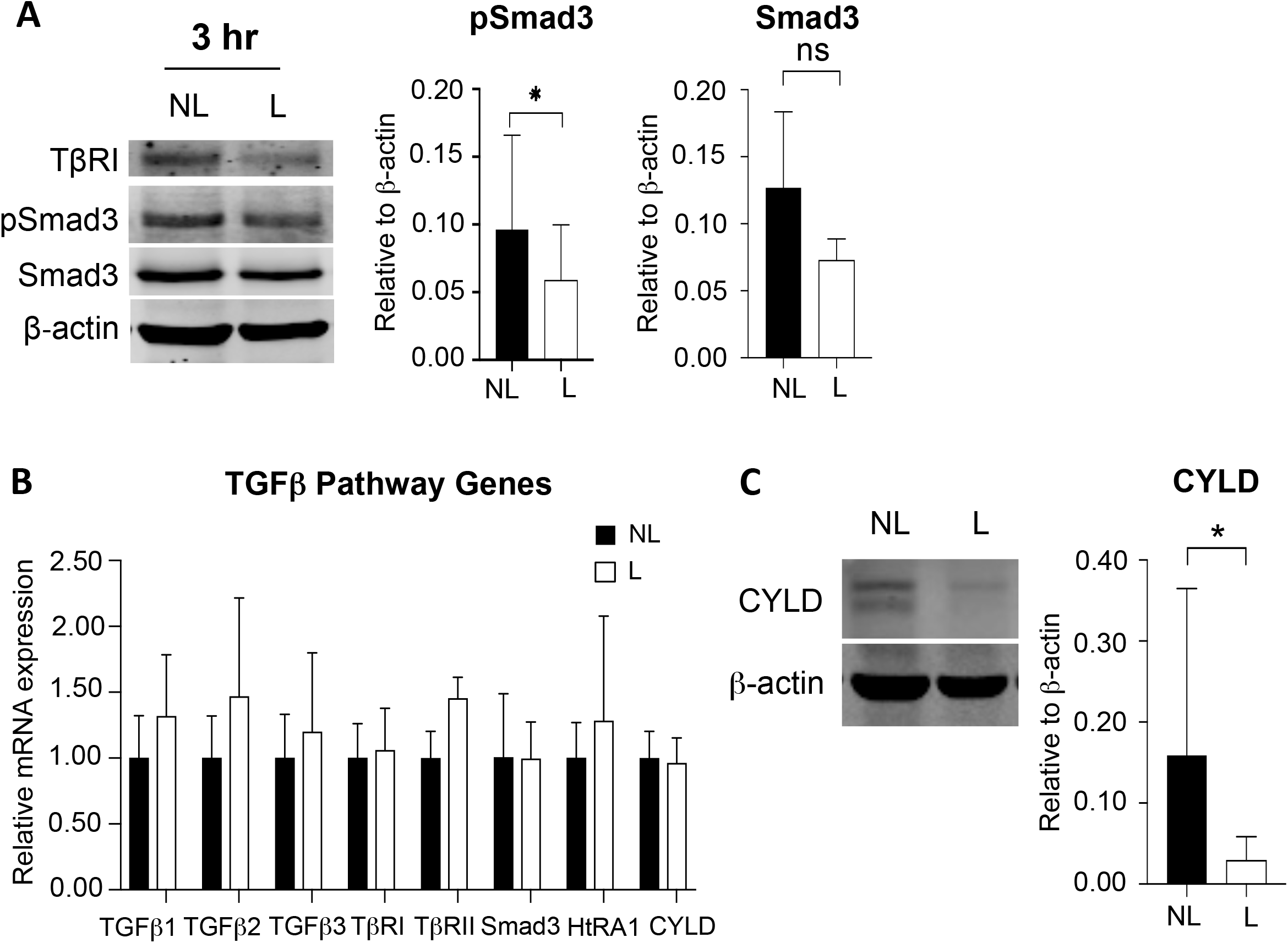
Rapid post-transcriptional control of TGFβ mechanosensitivity. (A) Protein levels of TGFβ receptor type 1 (TβRI), phosphorylated Smad3 (pSmad3), and Smad3 are reduced in loaded tibiae as compared to nonloaded tibiae (n>6). Phosphorylated Smad3 is reduced by 25% in loaded tibiae within 3hr of the mechanical stimulation. Error bars shown are standard deviation and the non-parametric Wilcoxin test was used for statistical analysis (n>6, *p<0.05). (B) mRNA levels for TGFβ ligands, receptors, Smad3, HtrA1, and CYLD are not significantly regulated by load within 3 hrs. Data shown are averages with standard error of the mean and the Wilcoxin test was used for statistical analysis (n>5). (C) CYLD protein levels are reduced in loaded tibia 3 hrs after stimulus. Error bars shown are standard deviation and the Wilcoxin test was used for statistical analysis (n>6, *p<0.05).

Interestingly, the expression of CYLD, a deubiquitinase implicated in TGFβ signaling in the lung [20], was significantly reduced by at least 50% within 3 hrs of loading (Figure 1C). These findings, as well as the absence of transcriptional differences in TGFβ signaling components at this time point, collectively focused our attention on post-transcriptional mechanisms responsible for TGFβ mechanosensitivity in bone.

### 3.2 Prostaglandins repress Smad3 transcriptional activity in a PGE2-dependent mechanism

To further dissect the mechanisms involved in the mechanoregulation of TGFβ signaling, we evaluated the regulation of TGFβ signaling by prostaglandins in MLO-Y4 osteocyte-like cells. PGE2 is an established early mediator of mechanotransduction and is rapidly increased within 10 minutes of fluid flow stimulation in osteocytes [26]. Blocking PGE2 signaling with a chemical antagonist of the EP2 receptor, AH6809, inhibits downstream effects of fluid flow stimulation in osteocytes [27]. Since PGE2 can repress TGFβ signaling in mammary tumor cells through EP2 [17], we hypothesized that PGE2 is sufficient to antagonize TGFβ signaling in osteocytes.

To test this hypothesis, we treated MLO-Y4 cells with 5μM PGE2 for 3 hrs. Following TGFβ stimulation, phosphorylated Smad3 activates transcription of TGFβ target genes such as Serpine1 (Figure 2A). However, PGE2 significantly reduces TGFβ-inducible Serpine1 mRNA expression (Figure 2A). PGE2 signals through EP2 and EP4 receptors. Butaprost, a pharmacologic agonist of the EP2 receptor, mimics the effects of PGE2 treatment by suppressing TGFβ-inducble Serpine1 mRNA expression (Figure 2B). The EP4 agonist, PGE1-alcohol, had similar effects (Figure 2C).

**Figure 2:**
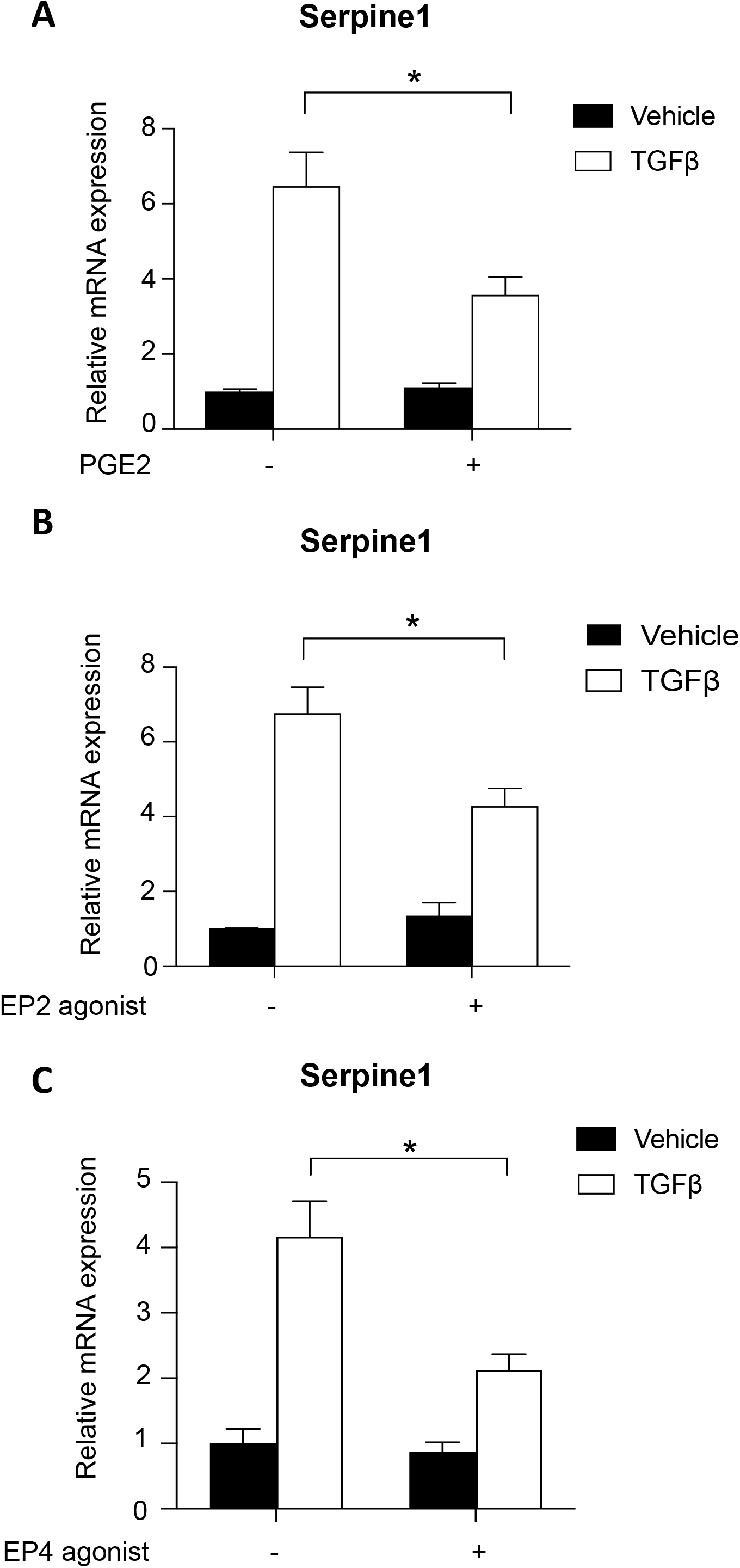
Prostaglandins repress Smad3 transcriptional activity in a PGE2-dependent mechanism. PGE2 inhibits the TGFβ responsive genes Serpine1 in MLOY4 cells within 3 hrs (A). Butaprost, an EP2 receptor agonist, also represses Serpine1 transactivation in MLOY4 cells within 3 hrs (B). Similarly, PGE1 alcohol, an EP4-agonist, also inhibits Serpine1 expression in MLOY4 cells within 3 hrs (C). All experiments were repeated at least three times, with two technical replicates for the RNA analysis. Standard error of the mean are shown (* p<0.05).

### 3.3 Prostaglandins suppress Smad3 phosphorylation through a proteasome-dependent mechanism

Since Smad3 phosphorylation and CYLD protein expression were regulated similarly in response to mechanical loading in vivo, we hypothesized that CYLD would also be regulated by prostaglandins in vitro. In the same conditions used to evaluate Smad3 phosphorylation, both PGE2 (Figure 3A) and the EP2 agonist (Figure 3B) reduced CYLD protein expression within 3 hrs. Therefore, prostaglandin signaling through EP2 receptors in cultured osteocytes is sufficient to repress Smad3 phosphorylation and CYLD expression in a manner that mimics the rapid repression in hindlimb-loaded bones.

**Figure 3:**
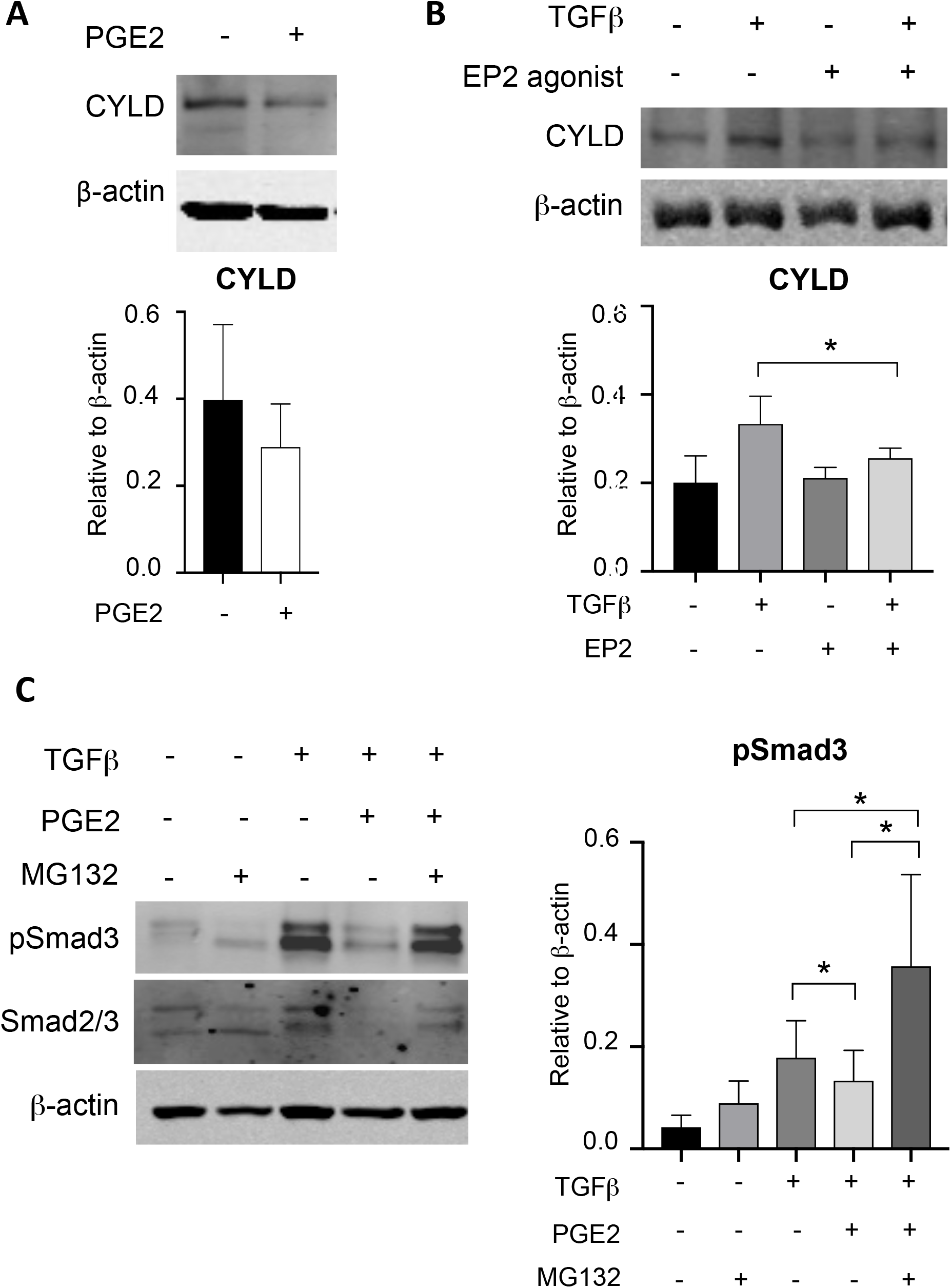
Prostaglandins suppress CYLD expression and Smad3 phosphorylation in a proteosome-dependent mechanism. (A) PGE2 represses CYLD protein expression in MLOY4 cells within 3 hrs. (B) Stimulation of EP2 with butaprost for 3 hrs in MLOY4 cells down-regulates CYLD protein expression. (C) MG132, a proteasome inhibitor, stabilizes TGFβ-activated pSmad3 levels during a 3 hrs exposure to PGE2 in MLOY4 cells. Data shown are averages of at least three experiments with standard deviation shown (* p<0.05).

The common regulation of Smad3 phosphorylation and CYLD expression by load in bone, and by prostaglandins in MLO-Y4 osteocyte-like cells, led us to investigate the epistatic interaction of Smad3 and CYLD. Since CYLD is a deubiquitinase that regulates proteolytic degradation [20], we tested the effect of proteosome inhibition on the regulation of Smad3 phosphorylation by TGFβ and prostaglandins. As in Figure 3, prostaglandins block TGFβ inducible Smad3 phosphorylation (Figure 3C). However, the inhibitory effect of prostaglandins on Smad3 phosphorylation is reversed in MLO-Y4 cells treated with the proteasome inhibitor MG132 (Figure 3C). This suggests that the repressive effect of prostaglandins on Smad3 phosphorylation in osteocytes relies on proteosomal activity.

### 3.4 CYLD is required for PGE2/EP2 repression of TGFβ signaling in osteocytes

To test the hypothesis that the deubiquitinase CYLD controls the level of Smad3 phosphorylation in osteocytes, we suppressed CYLD expression in MLO-Y4 cells. CYLD mRNA expression was 50% lower in MLO-Y4 cells transfected with CYLD siRNA, relative to cells transfected with a scrambled siRNA control (Figure 4A). Likewise, CYLD siRNA significantly reduced CYLD protein expression (Figure 4B) and increased the expression of CYLD-repressed genes, MMP2 and MCSF (Figure 4C) [28]. Using this approach, we found that CYLD is required for prostaglandin-dependent repression of Smad3 phosphorylation. As expected, the EP2 agonist significantly represses TGFβ-inducible Smad3 phosphorylation in control siRNA-transfected MLO-Y4 cells (Figure 4D). However, this repression is lost in cells transfected with CYLD siRNA. Likewise, the ability of prostaglandins to block TGFβ-inducible Serpine1 mRNA expression is abrogated in CYLD siRNA transfected cells (Figure 4E). Therefore, prostaglandins antagonize TGFβ signaling through Smad3 in a CYLD-mediated mechanism in osteocytes.

**Figure 4:**
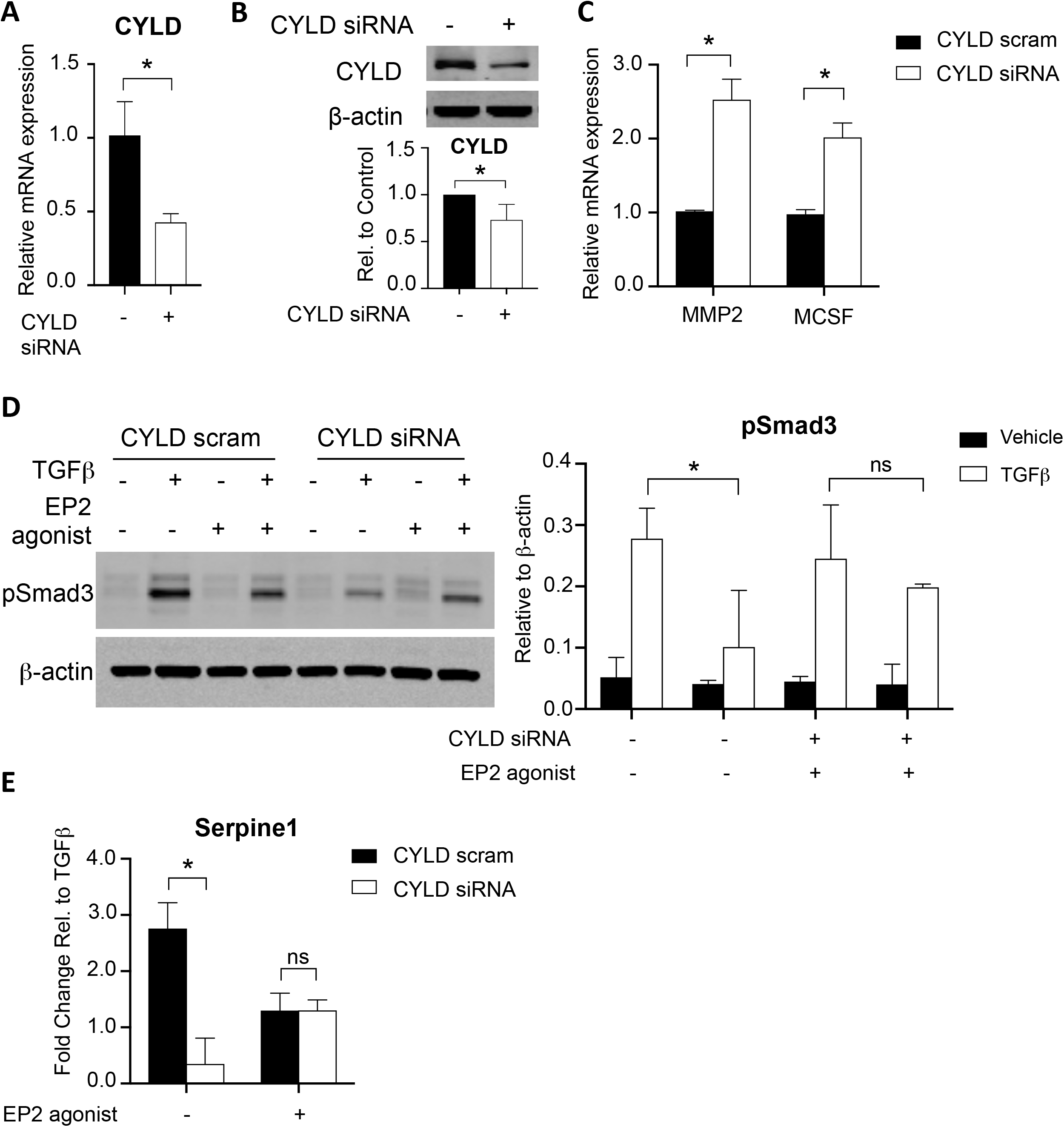
CYLD is required for PGE/EP2 repression of TGFβ activity in osteocytes. CYLD siRNA effectively reduced CYLD mRNA (A) and protein (B) expression relative to a scrambled control siRNA. CYLD protein expression is significantly reduced in MLOY4 cells containing CYLD siRNA (B). Relative to a scrambled control siRNA, siRNA for CYLD leads to increased expression of MMP2 and MCSF, known targets of CYLD repression (C). The EP2 agonist butaprost prevents TGFβ transactivation of pSmad3 in control siRNA MLOY4 cells, while CYLD siRNA rescues TGFβ-inducible Smad3 phosphorylation in the presence of EP2 stimulation (D). Likewise, reduced CYLD activity in CYLD siRNA MLOY4 cells prevents EP2-mediated repression of Serpine1 mRNA expression. This is a representative figure of at least three different experiments and average fold changes along with standard error of the mean are shown (E). All experiments were repeated at least three times and averages along with standard deviation (B, D) or standard error of the mean (A, C) are shown (p<0.05)

### 3.5 CYLD is required for load-mediated repression of pSmad3 and load-induced bone formation

The rapid epistatic control of Smad3 by prostaglandins and CYLD led us to hypothesize that CYLD participates in the mechanosensitive control of TGFβ signaling in bone anabolism. This hypothesis is consistent with the mechanosensitive repression of CYLD expression in loaded bone (Figure 1C). To test this hypothesis, we applied mechanical stimulation to tibiae of CYLD knockout mice and their wild type littermates [24]. As expected, within 3 hrs after mechanical load, loaded tibiae of wild type mice showed reduced levels of Smad3 phosphorylation relative to nonloaded contralateral tibia (Fig. 5A). In CYLD-deficient mice, however, loading caused no significant changes in the level of Smad3 phosphorylation (Figure 5A).

**Figure 5:**
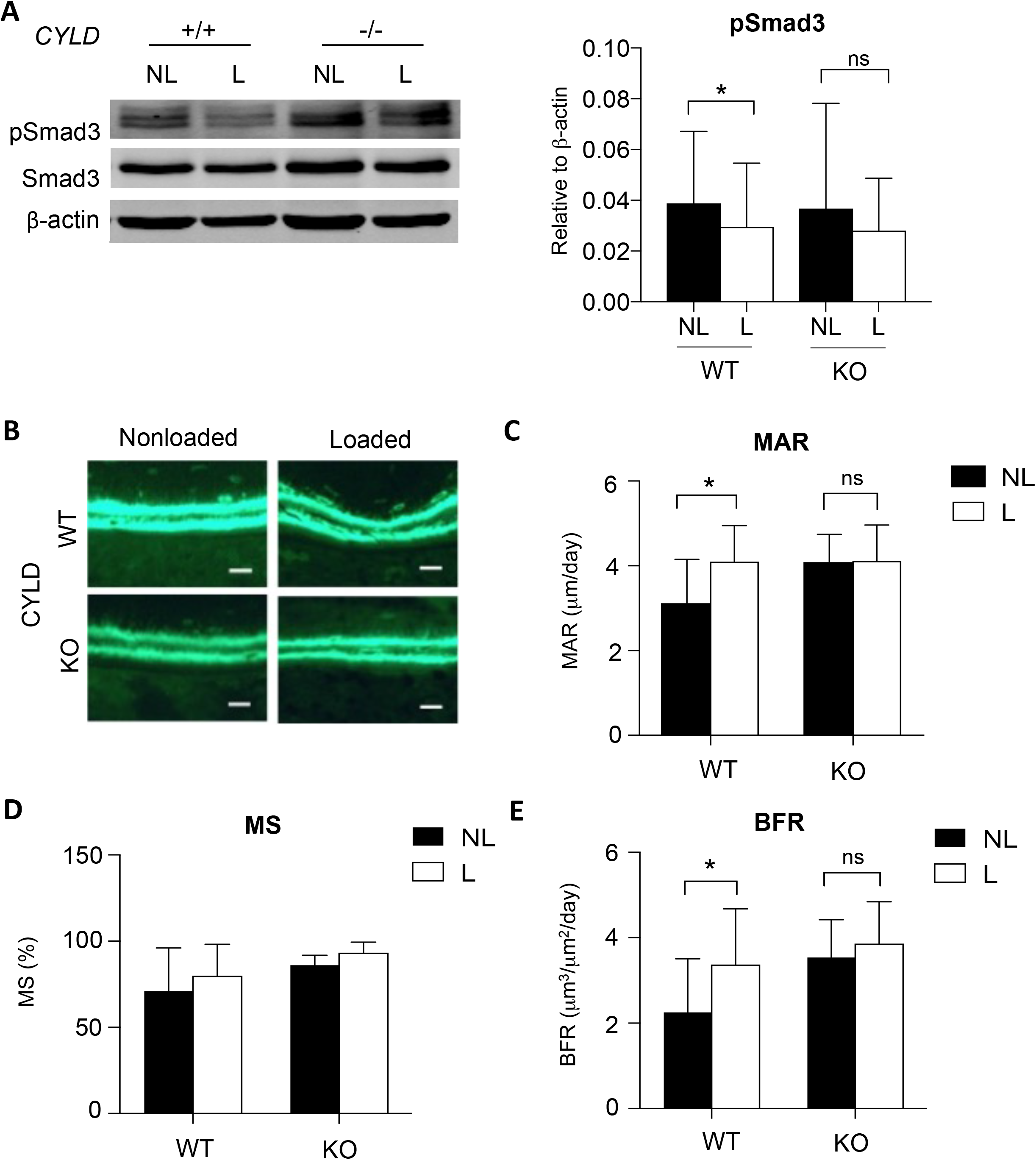
CYLD is required for load-mediated repression of pSmad3 and load-induced bone formation. pSmad3 activation is reduced in tibia from loaded (L) relative to non-loaded limbs from wild type (+/+) mice, but not from their CYLD-deficient (-/-) littermates (A). Data shown are averages along with standard deviation and the Wilcoxin test was used for statistical analysis (n>5, *p<0.05). Calcein labeling of newly formed bone shows a load-dependent increase in calcein incorporation in wild type mice, but not in CYLD-deficient littermates. Line indicates 20μM magnification. (B). Dynamic histomorphometry measurements of mineral apposition rate (MAR), mineralizing surfaces (MS), and bone formation rate (BFR) showed a significant increase in MAR and BFR in loaded tibiae of wild type mice, but not in loaded tibiae of their CYLD-deficient littermates (C-E). Data shown are averages along with standard deviation and paired t-test was used for statistical analysis (n>5, * p<0.05).

Dynamic histomorphometry was used to test the functional effects of CYLD-deficiency on load-induced bone formation in wild type and CYLD-deficient mice. Calcein labeling showed an increase in bone formation in loaded tibiae of wild type mice, relative to the non-loaded contralateral tibia (Figure 5B). However, no further increase was observed in loaded tibiae of CYLD knockout mice compared to control (Fig. 5B). Accordingly, the matrix apposition rate (MAR) and the bone formation rate (BFR) are significantly increased in loaded bone of wild type mice compared to control but not in loaded tibiae of CYLD knockout mice compared to control (Figure 5C and 5D). No significant load-dependent differences in the mineralization surface were observed in either wild type or CYLD-deficient bones (Figure 5E). Together these data indicate that CYLD is necessary for the rapid mechanosensitive repression of Smad3 phosphorylation, as well as for load-induced bone formation.

## 4. Discussion

Here we describe a new mechanosensitive mechanism that regulates TGFβ signaling in bone rapidly after hindlimb loading. TGFβ-inducible phosphorylated Smad3 was repressed within 3 hrs after mechanical loading on the tibia. We found that PGE2, acting through the EP2 receptor, is sufficient to repress pSmad3 and its transactivation of Serpine1 in osteocytes. PGE2 and EP2 suppress TGFβ signaling through a proteasomal mechanism that depends on the deubiquitinase CYLD. Furthermore, CYLD is mechanosensitive in bone and is required for load-induced bone formation (Figure 6). To our knowledge, this is the first study to uncover CYLD mechanosensitivity and to implicate CYLD as a target of the PGE2 pathway.

**Figure 6:**
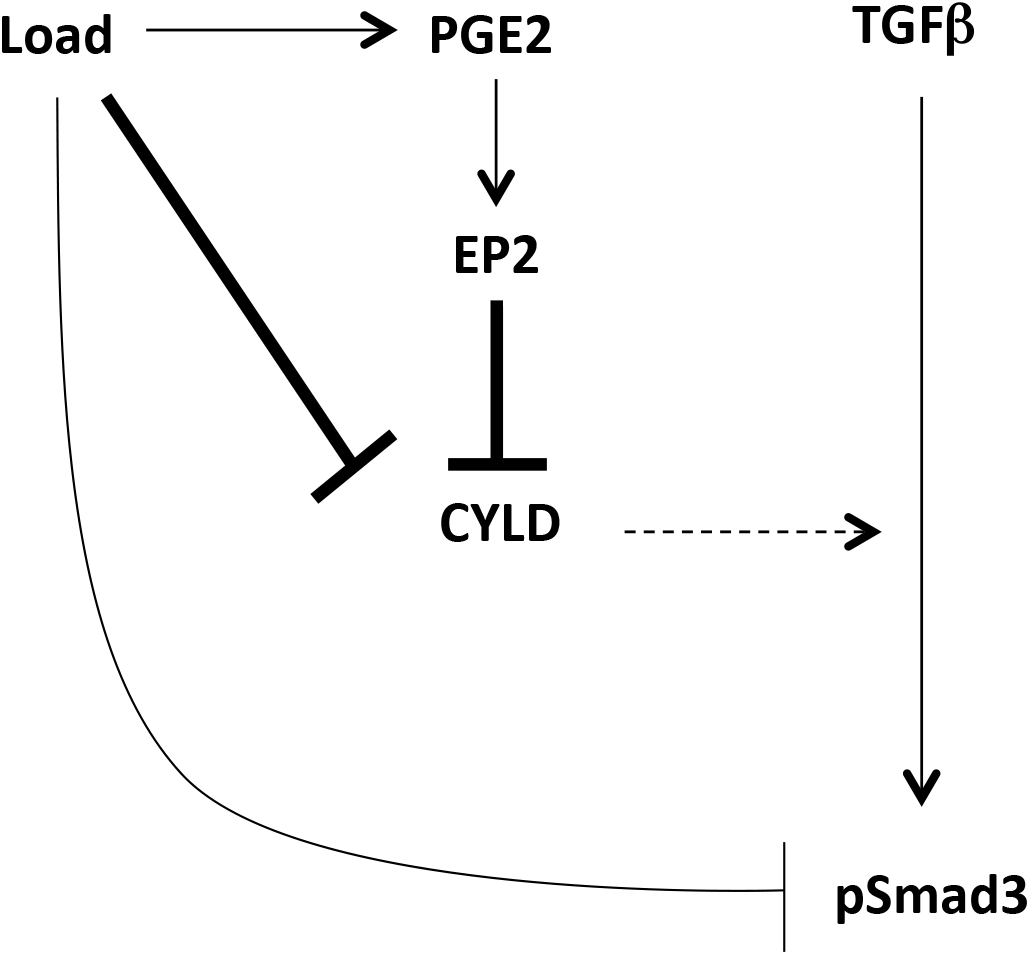
Mechanosensitive CYLD mediates PGE2 repression of TGFβ signaling. This study builds on our prior understanding of mechanisms by which load regulates TGFβ signaling in bone (light lines) by showing (bold lines) the sensitivity of CYLD expression to mechanical load and to prostaglandin signaling, and the requirement for CYLD in the regulation of Smad3 phosphorylation and load-induced bone formation.

TGFβ signaling is tightly regulated by many physical cues, including compression, tension, substrate stiffness, and fluid flow shear stress [8] [29] [30] [31]. This regulation occurs at multiple levels of the TGFβ pathway. In some cases, the same physical stimulus simultaneously exerts distinct and competing effects on TGFβ ligands, receptors, and intracellular effectors. For example, after 6h of fluid flow shear stress, renal epithelial cells produce lower levels of total and active TGFβ1 ligand, but possess elevated levels of TGFβ1 mRNA and increased levels of phosphorylated Smad2/3 [32]. On the other hand, interstitial fluid flow on dermal fibroblasts for 2 hrs increases TGFβ1 secretion but also decreases TβRII protein expression [33]. With respect to bone, periosteal cells from rat hindlimbs possess increased levels of TGFβ1 mRNA expression within 4 hrs of 4-point bending [34]. In tibiae of loaded wild type mice, we find that TGFβ ligand mRNA expression is not significantly altered, although pSmad3 is reduced 3 hrs after load. Consistent with evidence throughout the field of mechanobiology, the effect of physical cues on TGFβ signaling depends on the cell type, the physical perturbation, the time points, and the cellular microenvironment, among other factors. However, the molecular mechanisms responsible for the diversity of these responses remain unclear. Within this context, we implicate a novel CYLD-dependent mechanism by which cells modulate TGFβ signaling in response to physical cues.

PGE2 is vital to maintaining bone homeostasis and so it functions in many biological processes including mechanotransduction. This inflammatory mediator signals through four different G-protein receptors, EP1-4, with active EP2 and EP4 inducing bone formation [35] [36]. Inhibiting PGE2 synthesis with Naproxen, an NSAID, treatment in mice impairs load-induced bone formation and reduces NGF expression [37]. EP4 functions downstream of PGE2 to regulate processes for bone homeostasis including crosstalk between the nervous system and bone formation [38]. Accordingly, the EP4 agonist, ONO4819, additively enhances load-mediated bone formation in rats [39]. In osteocytes, the cells that modulate the mechanoresponse in bone, fluid flow stimulation induces PGE2 and EP2 production [27] [14]. Since EP2 and TGFβ are both mechanosensitive in osteocytes, we hypothesized that EP2 may regulate TGFβ function in these cells [8]. We show here that PGE2 and EP2 repress TGFβ signaling in osteocytes through a proteasome-mediated mechanism that depends on the activity of the deubiquitinase CYLD. Future in vitro studies with fluid flow shear stress will help to dissect the molecular epistasis of PGE2, EP2, CYLD, and TGFβ signaling in osteocyte mechanotransduction.

CYLD and TGFβ signaling intersect at many levels. CYLD regulates the stability of TβRI, Smad7, and Smad3 [40] [20] [28]. In bone, CYLD inhibits osteoclastogenesis. With hyperresponsivity to RANKL, CYLD-deficient mice thus exhibit low trabecular bone volume, but no significant change in cortical bone volume [23]. We found that CYLD is repressed with mechanical stimulation and is important for load-mediated bone formation. This introduces CYLD as one of the few deubiquitinases to be mechanosensitive, including CHIP and MEC-4 [18] [19]. While we uncovered the role of CYLD in regulating TGFβ signaling in mechanically stimulated osteocytes, CYLD regulates effectors of other pathways such as the Wnt pathway [41]. Studying how CYLD intersects with other mechanosensitive pathways can uncover how signaling is dampened and controlled for proper homeostasis. Further study of the role of CYLD in bone mechanobiology may reveal CYLD to be an attractive therapeutic target for osteoporosis.

## Acknowledgements

Support of this study was provided for JN by the NIH through F30 DE022680 and for TA by the NIH through R01 DE019284 and R21 AR070403, the NSF through NSF 1636331, and the DOD through OR130191 and OR170044.

